# CimpleG: Finding simple CpG methylation signatures

**DOI:** 10.1101/2022.09.12.507513

**Authors:** Tiago Maié, Marco Schmidt, Myriam Erz, Wolfgang Wagner, Ivan G. Costa

## Abstract

DNA methylation (DNAm) at specific CG dinucleotides (CpG sites) in the genome provides powerful biomarkers for a wide range of applications, such as epigenetic clocks, cell type deconvolution, and stratification of cancer patients. Such epigenetic signatures are usually based on multivariate approaches that require hundreds of DNAm sites for predictions. In contrast, targeted assays for only a few selected CpGs may facilitate faster turnover, better standardisation, and reduced costs. Here, we propose a computational framework named CimpleG for the detection of small CpG methylation signatures, exemplarily used for cell type classification. We evaluate the performance of CimpleG and competing methods on two distinct datasets in regard to the cell-type classification of blood cells and of other somatic cells. Furthermore, we evaluate how well these classifiers perform cellular deconvolution of blood cell mixtures. We show that CimpleG is both time efficient and performs as well as Elastic Net, while basing its prediction on a single DNAm site per cell type. Altogether, CimpleG provides a complete computational framework for the delineation of DNAm signatures and cellular deconvolution.

## Introduction

DNA methylation (DNAm) is intrinsically related to chromatin structure and cell differentiation and controls the expression of neighbouring genes^1^. The stability of DNAm and the accuracy of genomic methods for quantification of DNA methylation levels makes DNAm a powerful molecular marker for the prediction of age^2–5^ and stratification of cancer patients^6, 7^. Furthermore, some loci have a characteristic DNAm profile in specific cells and can therefore be used for cell type characterisation^8, 9^ and estimation of cell proportions in tissues via cellular deconvolution^10, 11^. Development of such biomarkers was particularly eased by the rapidly growing number of available DNAm profiles that were measured with Illumina BeadChips, which can address up to 850.000 CpG sites^12^. The epigenetic signatures that are generated with these genomic DNAm profiles often comprise more than 100 CpGs. While the integration of multiple CpGs is considered to increase precision, it greatly hampers application in clinical settings, that require fast, standardised, and cost-effective analysis^13^. To this end, targeted analysis of individual CpGs e.g., with pyrosequencing, digital droplet PCR, epiTYPER, or amplicon sequencing may be advantageous^14^.

From a computational perspective, Elastic Net^15, 16^ is a commonly used method for building DNAm-based models, due to its ability to cope with large dimensional data by the selection of active features during model estimation. For example, a frequently used epigenetic ageing clock of Horvath is based on only 346 CpG sites^2^ selected with Elastic Net, from DNAm data measured with Illumina BeadChips (27k and 450k). Deep learning approaches displayed high predictive accuracy in cancer prediction, as recently shown in the prediction of primary or metastasis lung cancers^17^. These models, however, work on previously selected DNAm panels with only a small proportion of the initial DNAm sites (∼ 2000 DNAm sites) and do not indicate the importance of individual DNAm sites. A common approach for the detection of DNAm signatures for tissue deconvolution are statistical tests, which characterise differential methylated cytosines (DMC) between samples in two biological conditions^11^. Such approaches are provided by pipelines for analysis of DNAm^18–21^ and are mostly based on the use of the limma moderated *t*-test^22^ or on fold change statistics^23^. Altogether, we are not aware of any computational approach tailored for the selection of small DNAm signatures for cell type prediction and deconvolution.

We have recently described studies on the use of few CpG sites for cell-type deconvolution^13, 24^. Among others, we showed that DNAm signatures with two DNAm sites for fibroblast cells, which we used as a surrogate for fibrosis level, indicated lower survival rates in several cancer types. Building upon this work, we propose and formalise a computational framework named CimpleG for the detection of simple (small) CpG methylation signatures (Fig. 1A). In brief, CimpleG uses a univariate feature selection by combining a *t*-statistic measure with the Area under the Precision-Recall Curve (AUPR)^25^ to select the best DNAm sites for cell type classification. We evaluate CimpleG and competing methods in their performance on cellular prediction on two cell specific datasets with distinct types of blood and other somatic cells in both cell type prediction and cell deconvolution problems.

**Fig. 1.**
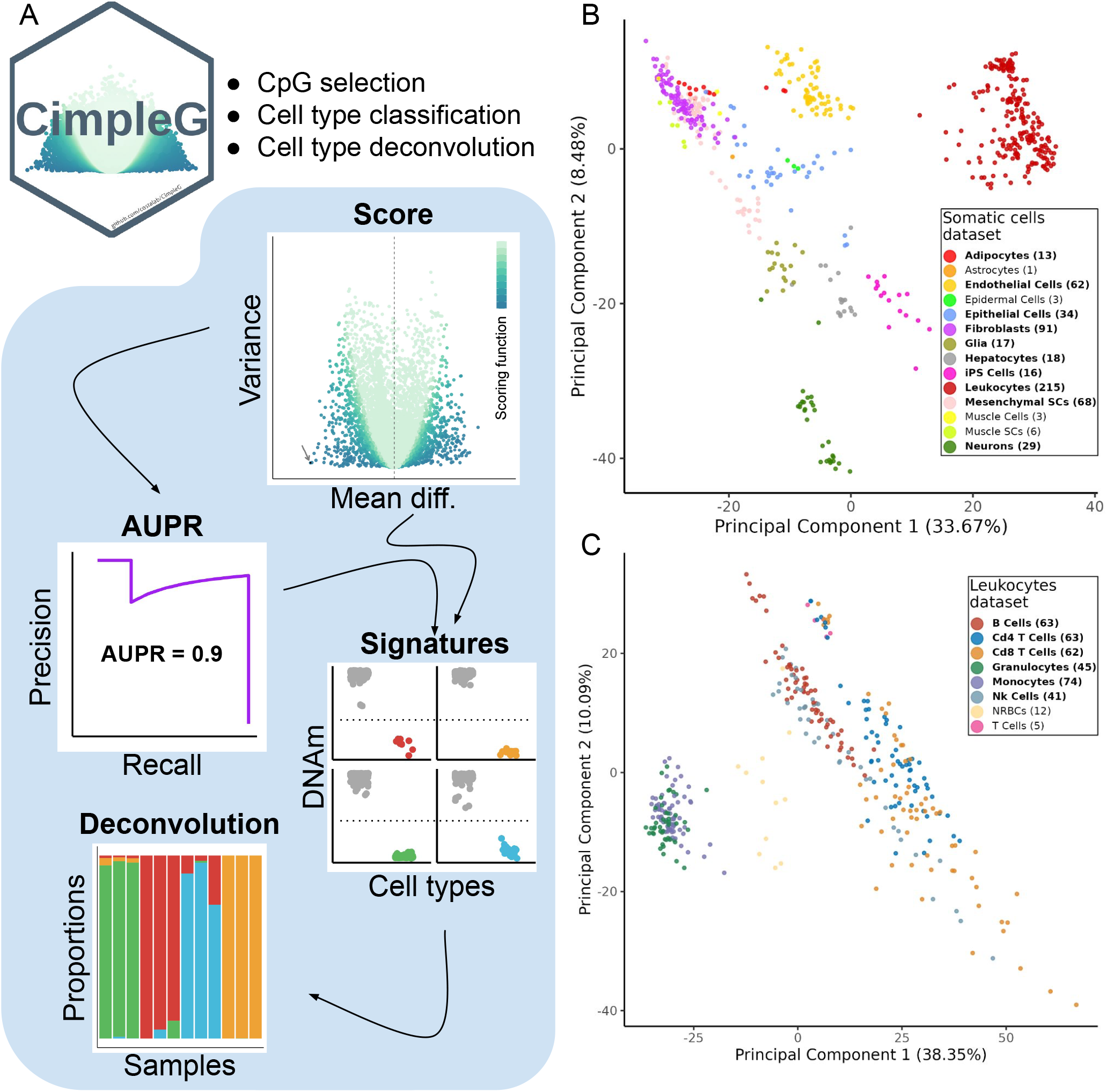
(A) Overview of CimpleG and statistics used for feature selection and downstream applications. (B-C) Principal component analysis of the DNAm datasets with somatic cells (B) or leukocytes (C). Only cell types highlighted in bold, for which we have samples in train (10 positive examples) and test data, were used as target classes. Cell types that are not present in the test data are only used as negative examples (non-target cells).

## Results

### CimpleG Framework

CimpleG is a computational framework for the selection of DNAm signatures for cell type classification (Fig. 1A). It provides a novel feature selection metric to select small DNAm signatures. CimpleG initially uses a *t*-statistic score to pre-select active features followed by the area under the precision-recall curve (AUPR) for feature selection. The use of precision-recall curves instead of the usual ROC curves for feature selection^25^ was adopted due to the high imbalance of classes in DNAm cell classification problems, i.e. an average of 15 negatives for a positive example. Next, CimpleG ranks and selects the top candidate CpG sites by combining the score and the AUPR value. These are used to build univariate cell-type-specific classifiers and for cell-deconvolution (Fig. 1A). In addition, the CimpleG framework facilitates the use of alternative feature selection and classification methods (such as random forests^26^, Elastic Net^16^ and Boosted trees (XGBoost)^27^).

Moreover, CimpleG provides two curated and pre-processed DNAm datasets with a compendium of DNAm arrays with 14 somatic cell types and eight different leukocytes (Fig. 1B-C). These data were pre-processed with well-known state-of-art DNAm-based methods such as SeSAMe^21^, minfi^18^ and watermelon^28^. The final somatic cells and leukocytes datasets have 576 and 365 samples with 143,291 and 284,706 CpG sites respectively. We stratified these datasets in train and test samples such that data from the same study are only found as test or training data. Moreover, training and test data were pre-processed independently to avoid leak pre-processing^29^ (see Table 1 and Table 2 for data characteristics). Principal component analysis of these two datasets shows some separation between major cell types (Fig. 1B-C), while closely related cells (Fibroblast and MSC cells in somatic cell data; CD4 and CD8 T cells in the leukocytes data) can only be discriminated with additional PCs (Supplementary Fig.S1).

**Table 1.** Somatic cells data summary.

|  | Train dataset | Test dataset | *Total* |
| --- | --- | --- | --- |
| **Adipocytes** | 10 (2.4%) | 3 (1.8%) | 13 (2.3%) |
| Astrocytes | 1 (0.2%) | 0 (0%) | 1 (0.2%) |
| **Endothelial Cells** | 32 (7.8%) | 30 (18%) | 62 (11%) |
| Epidermal Cells | 3 (0.7%) | 0 (0%) | 3 (0.5%) |
| **Epithelial Cells** | 16 (3.9%) | 18 (11%) | 34 (5.9%) |
| **Fibroblasts** | 52 (13%) | 39 (24%) | 91 (16%) |
| **Glia** | 10 (2.4%) | 7 (4.2%) | 17 (3.0%) |
| **Hepatocytes** | 15 (3.6%) | 3 (1.8%) | 18 (3.1%) |
| **iPS Cells** | 13 (3.2%) | 3 (1.8%) | 16 (2.8%) |
| **Leukocytes** | 182 (44%) | 33 (20%) | 215 (37%) |
| **Mesenchymal SCs** | 56 (14%) | 12 (7.3%) | 68 (12%) |
| Muscle Cells | 3 (0.7%) | 0 (0%) | 3 (0.5%) |
| Muscle SCs | 0 (0%) | 6 (3.6%) | 6 (1.0%) |
| **Neurons** | 18 (4.4%) | 11 (6.7%) | 29 (5.0%) |
| *Total* | 411 (100%) | 165 (100%) | 576 (100%) |

The highlighted rows indicate the cell types that were used to train the classifier.
The other cell types were only part of the training or test dataset for additional control.

**Table 2.** Leukocyte data summary.

|  | Train dataset | Test dataset | *Total* |
| --- | --- | --- | --- |
| **B Cells** | 50 (19%) | 13 (12%) | 63 (17%) |
| **CD4 T Cells** | 50 (19%) | 13 (12%) | 63 (17%) |
| **CD8 T Cells** | 50 (19%) | 12 (12%) | 62 (17%) |
| **Granulocytes** | 32 (12%) | 13 (12%) | 45 (12%) |
| **Monocytes** | 50 (19%) | 24 (23%) | 74 (20%) |
| **Nk Cells** | 29 (11%) | 12 (12%) | 41 (11%) |
| NRBCs | 0 (0%) | 12 (12%) | 12 (3.3%) |
| T Cells | 0 (0%) | 5 (4.8%) | 5 (1.4%) |
| *Total* | 261 (100%) | 104 (100%) | 365 (100%) |

The highlighted rows indicate the cell types that were used to train the classifier.
The other cell types were only part of the training or test dataset for additional control.
NRBCs = nucleated red blood cells

### Benchmarking of CimpleG and classification methods

CimpleG was evaluated in comparison to different alternative methods to generate epigenetic signatures: decision trees, random forests, boosted trees, neural networks and Elastic Net. We also considered a single-feature DNAm brute force classifier as a baseline. This brute force algorithm evaluates all possible individual markers and ranks these by AUPR. Moreover, we also evaluated CimpleG variants only considering the AUPR (CimpleG AUPR) or the t-statistic (CimpleG Score). This was to ensure that the combination of these metrics, was stronger than their individual use. Notably, some models (neural networks, random forests and decision trees) could not cope with the high dimensional feature size of the datasets. This was mainly due to very large memory usage or days of execution time. Therefore, for these models, as a pre-training step, we have performed an unsupervised feature selection considering variance and co-variance of the DNAm sites^30^. Next, we have used a cross-validation framework to optimise parameters for all methods. We evaluated three main benchmarking metrics, the classification performance as measured by the AUPR, the computational time required and the number of features used by each model. We only build classifiers/signatures for cell types with at least 10 samples in the training data (10 out of 14 in the somatic cells and 6 out of 8 in blood cells), however, we still keep cells with small sample sizes to be used as negative examples (non-target classes). See Supplementary Fig. S2 for an overview of the experimental pipeline.

We observed that Elastic Net, CimpleG and CimpleG (score) have the highest median AUPR for the somatic cells and the leukocyte dataset (Fig. 2A-B), indicating that these are the best performing models. By considering the means of ranks, we observed that Elastic Net, CimpleG and CimpleG (score) are the three best classifiers in regards to their accuracy on the test datasets. Since the number of target samples differs per target cell type, a relevant question is if there is any association between the number of positive examples (target samples) and the classifier’s accuracy of individual methods (Supplementary Fig. S3). We observed that the top-performing methods (Elastic Net and CimpleG) have stable AUPR values across positive sample numbers, which indicates they are robust regarding a low number of positive samples. Regarding computational time, per signature, CimpleG took on average 55.3 seconds, Elastic Net needed on average 37.6 minutes, while the Brute force algorithm required on average 6.61 hours to generate a signature (Fig. 2C-D). These results indicate these three methods perform equally well in the DNAm-based cell classification problem, while CimpleG provides a significant speed up for the feature selection problem.

**Fig. 2.**
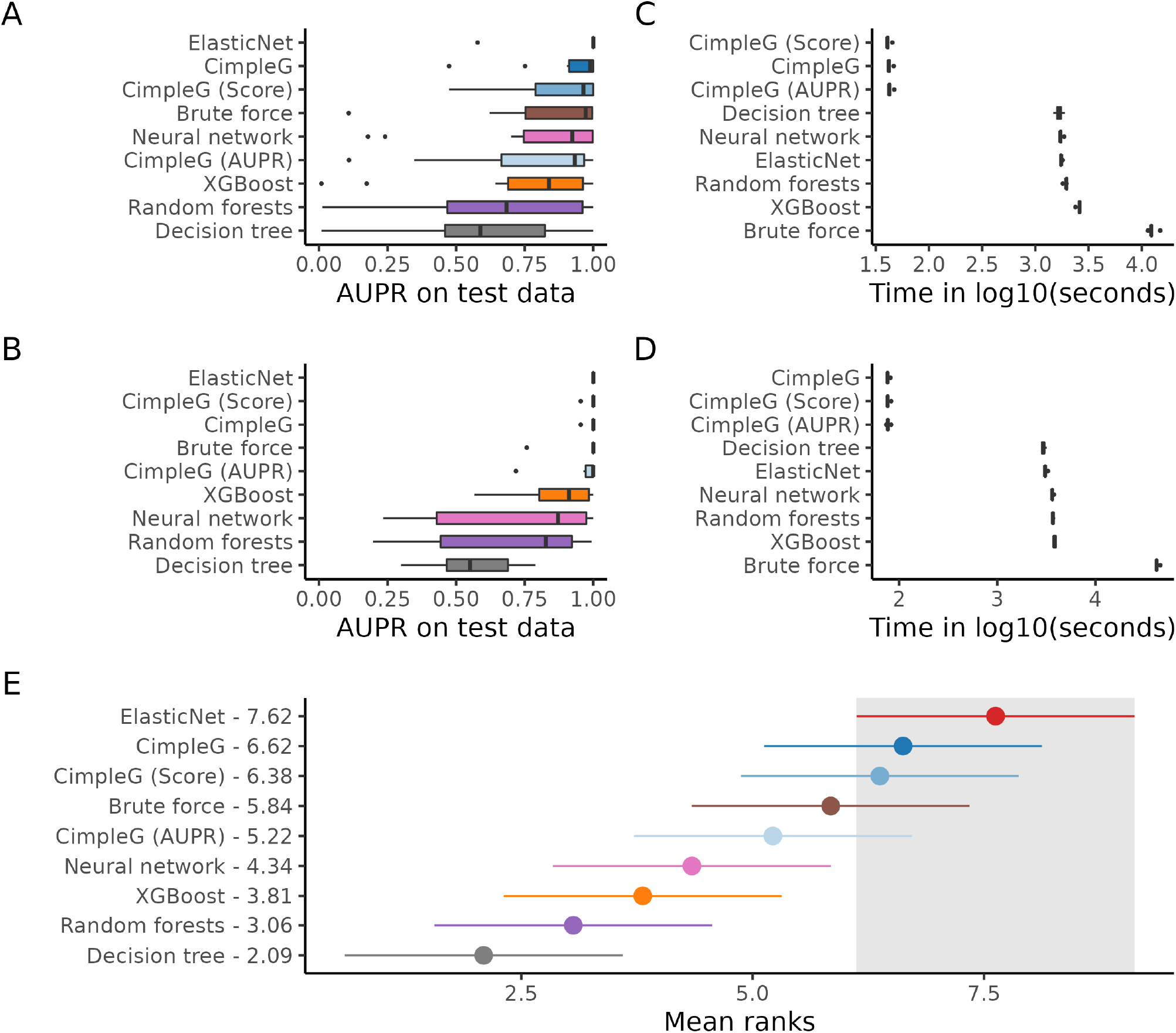
(A-B) Classification performance (AUPR) on the test set of the somatic cells (A) and leukocytes datasets (B). (C-D) Computational time required for each method to produce a signature including cross-validation and testing for the somatic cells (C) and leukocytes (D) datasets. (E) Mean ranks of the methods across all datasets based on the AUPR. The best method, Elastic Net, and its 95% confidence interval (Friedman and Nemenyi post-hoc test) is highlighted in grey. Methods that do not overlap at all with the highlighted area, are significantly worse than the highlighted methods.

### Selection of DNA methylation sites

Another relevant point is the number of CpGs that are implemented in the signatures derived by the different methods. Elastic Net selected the largest number of features with 3378 unique features across all six models for the leukocyte cells (Fig. 3A) and 2345 unique features across all ten models for the somatic cells (Supplementary Fig. S4A). This is more than all other models combined. The single-feature classifiers (CimpleG and Brute force) selected in total 10 and 6 DNAm sites for somatic and leukocytes data respectively, one CpG per cell type. We observed a high overlap of these features with the ones selected by the Elastic Net, i.e. all 16 DNAm selected by CimpleG were also part of the DNAm sites selected by Elastic Net. Interestingly, features selected by random forests, neural networks or decision trees were quite distinct from other methods. This rose from the fact that these methods require the previous use of univariate filters, due to their incapacity to deal with large dimensional inputs. Altogether these results show the ability of CimpleG in the delineation of small DNAm signatures.

**Fig. 3.**
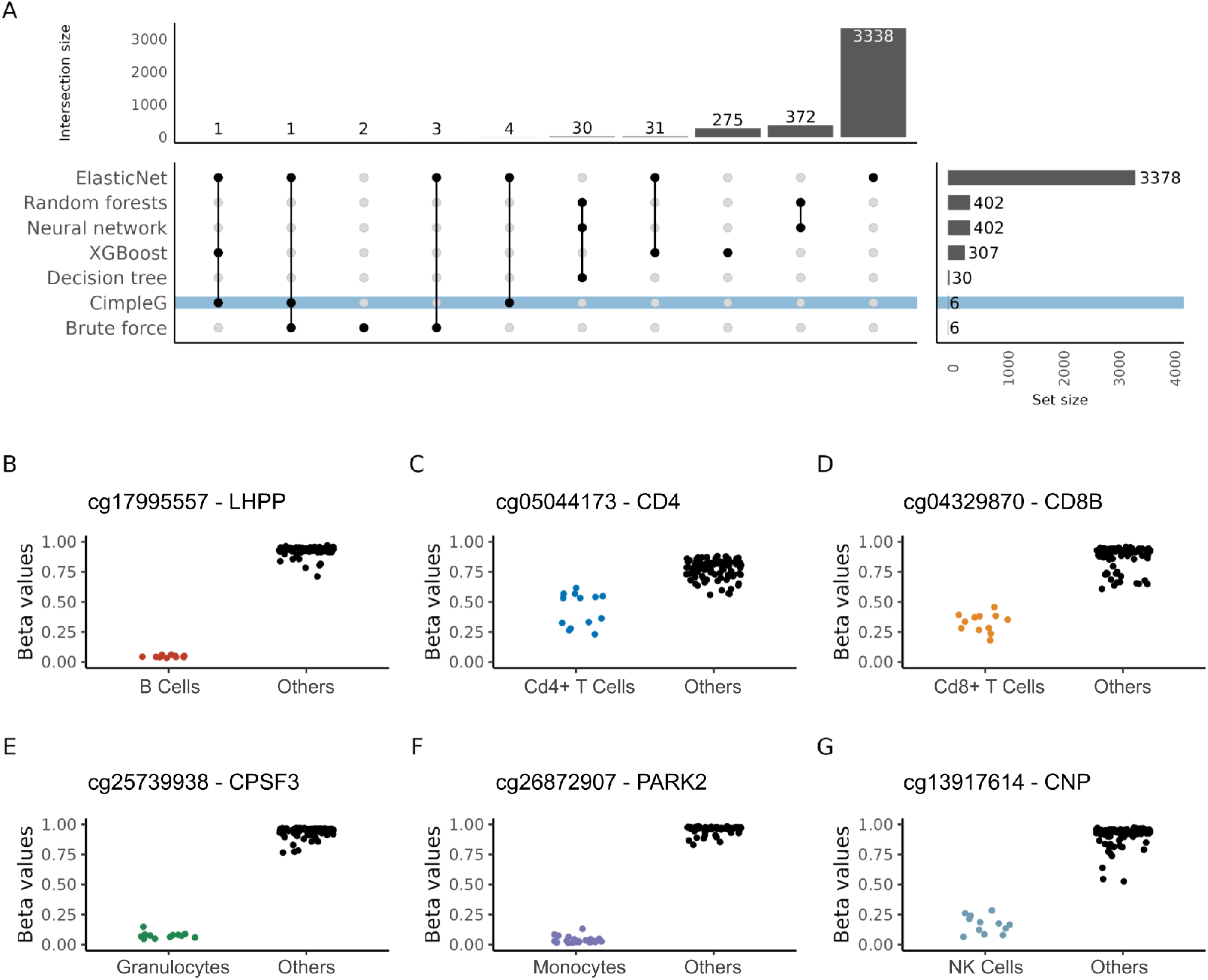
(A) Upset plot showing the total number of selected DNAm sites per method (y-axis) and how these are shared per method combinations (x-axis) for the leukocytes dataset. Connected dots in a column indicate the combination of methods considered in the x-axis. (B-G) Beta values of CpG sites selected by CimpleG on the test data. The colour of the points corresponds to the target cell type, while points in black correspond to the cell types that are not the targets for that signature.

Furthermore, it is interesting to look at the specific signatures generated by CimpleG (Fig. 3B-G; Supplementary Fig. S4B-K) as these genomic locations could provide biological insight into the cells themselves (see Supplementary Table 1 for complete results). Some DNAm sites are close to genes functionally related to cells, i.e. DNAm sites close to genes for CD4 (cg05044173, Fig. 3C) and CD8 (cg04329870, Fig. 3D) are selected as markers for CD4+ and CD8+ T cells. A DNAm site (cg01537765, Supplementary Fig. S4B) in the body of LIPE (Lipase E, Hormone Sensitive Type) is selected as a marker for adipose cells. This gene is known to function in adipose tissue by hydrolysing stored triglycerides to free fatty acids^31^. Finally, a CpG (cg10624122, Supplementary Fig. S4J) in the promoter of the epithelial mesenchymal transition related transcription factor, TWIST1^32^, is selected as a marker for mesenchymal stem cells. Moreover, other CpGs are close to genes with cell-specific expression patterns according to the human protein atlas^33^. An example is the CpG cg10673833 (Supplementary Fig. S4I) close to the gene MYO1G, which is a good marker for lymphocytes; and the CpG cg23882131 (Supplementary Fig. S4E) close to MRGPRF, which is a marker for fibroblasts. While the functional association of other markers is not evident, it needs to be considered that DNAm status does not generally translate directly to the expression of neighbouring genes.

### Deconvolution

Next, we evaluated the DNAm signatures and model predictions on a cell deconvolution problem with leukocytes. For this, we use either the DNAm sites (for models with small signatures; CimpleG, Brute Force) or the model prediction scores (for Elastic Net, Random Forests, Boosted Trees and Neural Networks) to build reference matrices for each model vs. cell type. We used these as input for the deconvolution method, the non-negative least squares (NNLS) algorithm^34^, which is a common deconvolution method for DNAm due to its simplicity and being among the top performers in a recent benchmarking study^35^. Methods were evaluated based on predictions on artificial blood cell mixtures of samples that were recently reported by Salas et al.^36^. From the models tested, we observe that CimpleG performs the best both in terms of the error associated with the prediction (RMSE) and fit compared to expected values (R^2^) (Fig. 4A-B). Excluding CimpleG, Elastic Net and Brute force, whose cell-specific predictions are shown in Fig. 4E-G, other models do not seem to perform well on the deconvolution task (Fig. 4C-D) as can be seen by their high RMSE and low R^2^ values (Supplementary Fig. S5). These results support the power of small signatures with respect to the problem of cellular deconvolution.

**Fig. 4.**
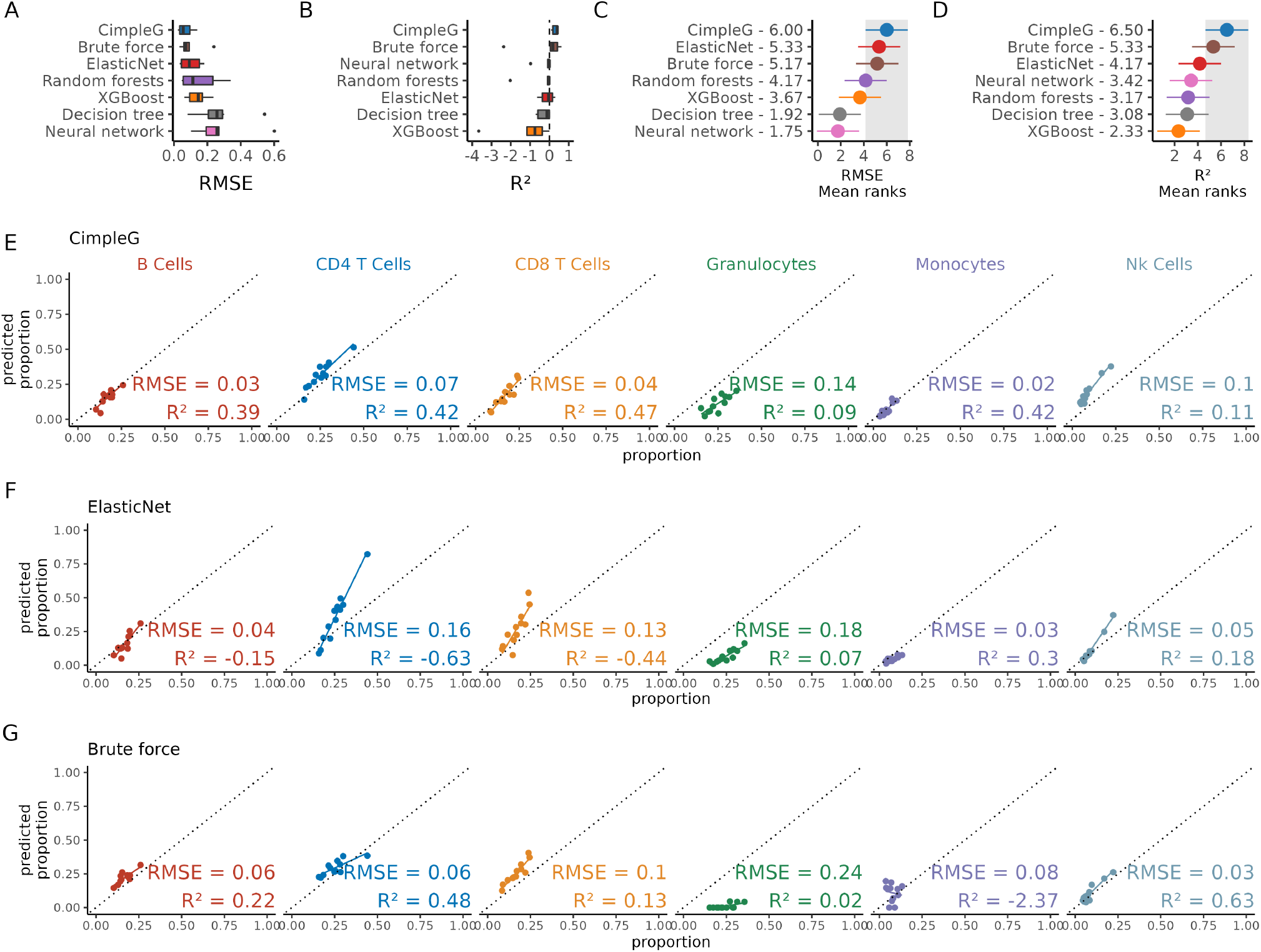
(A-B) Performance of models on the deconvolution on a artificial mixture data as measures by (A) RMSE (lowest is best) and (B) R^2^ (highest is best). (C-D) Overall mean ranks of the methods based on the RMSE and R^2^ respectively. The best method, CimpleG, and its 95% confidence interval (Friedman and Nemenyi post-hoc test) is highlighted in grey. Methods that do not overlap at all with the highlighted area, are significantly worse than the highlighted method. (E-G) The predicted proportion (y-axis) vs real proportion (x-axis) for all leukocyte mixtures for the top three performing methods: CimpleG (E); Elastic Net (F) and Brute force (G). Predictive accuracy statistics, RMSE and R^2^, are shown for each trained classifier.

## Discussion

Despite a huge number of DNAm biomarkers based on large CpG signatures, hardly any of them have been translated to clinical practice^14^. Many of the epigenetic signatures comprise a multitude of CpGs, which requires microarray or deep-sequencing methods that are difficult to implement for routine applications. Furthermore, even bioinformatic pipelines that need to be applied during analysis have to be certified according to the regulatory guidelines e.g., for in vitro diagnostic devices. In contrast, DNAm at individual CpG sites can be measured with a wide range of alternative methods, such as pyrosequencing, ddPCR, epiTYPER, or amplicon bisulfite sequencing^37, 38^. To this end, a framework is required that can identify the most powerful selection of individual CpGs for a given application.

We propose here CimpleG, which explores both t-statistic and AUPR scores, to select a single DNAm site per cell type of interest. The performance of CimpleG was contrasted to other state-of-art methods, such as Elastic Net, Random Forests, Boosted Trees and Neural Networks. Our results indicate that Elastic Net, which is broadly used for ageing signatures^2^, is the overall best method, but tends to select relatively large signatures that can comprise thousands of DNAm sites for its predictions. CimpleG performed as accurately as Elastic Net while focusing on only one DNAm site per cell type. Finally, we evaluated the performance of all models in the deconvolution of artificial mixture data. Here, CimpleG outperformed all competing methods, while Elastic Net was ranked second. It might be counter-intuitive that the simple single CpG signatures outperformed the larger signatures in the deconvolution of cell types. One reason is that large signatures are more prone to missing values for individual hybridisations.

Altogether, the CimpleG framework can be seen as a general pipeline for DNAm signature selection. It is implemented in R and provides functionalities to estimate both small DNAm signatures (using CimpleG), but also to derive Elastic Net and other machine learning methods evaluated here. It provides two large, manually curated and normalised datasets for the testing and benchmarking of methods. Moreover, cell type classifiers can be integrated with the NNLS method for cellular deconvolution, which makes it the first end-to-end pipeline for the estimation of cell-specific small DNAm signatures to cellular deconvolution. We are not aware of any other computational package addressing these.

There are further aspects to be explored in the future, such as synergistic effects of the small DNAm signatures on distinct deconvolution methods such as NMF^39^ and EpiDISH^23^; the impact of missing reference cell types or the use of CimpleG signatures for cell-of-origin detection from circulating cell free DNA^40^. From a technical perspective, there is a need for data structures for the efficient handling of the ever-increasing DNAm datasets. Another relevant issue are approaches for automatic integration with data repositories such as Gene Expression Omnibus. Note, however, that the lack of a consistent cell type annotation, i.e. as provided in cell ontology^41^, on such repositories, makes manual cell type annotation still a requirement. Taken together, CimpleG provides a valuable tool for clinician scientists as an easy-to-use framework for the identification of CpG sites that are best suited as a biomarker for targeted DNAm analysis.

## Methods

### CimpleG

CimpleG receives as input a matrix **X** ∈ (0, 1) ⊂ ℝ*p*×*n*;, where *x*_*ij*_ represents the DNAm values for site *i* in observation *j, n* is the number of observations and *p* is the number of DNAm sites. This essentially describes a matrix of Beta-values. The Beta-value is the ratio of methylated probe intensity versus the overall intensity in the Illumina methylation assays ^42, 43^. Alternatively, CimpleG can also, receive as input a matrix of M-values, these are the log2 ratio of intensities of methylated versus unmethylated probes^42^, a metric widely used in microarray assays and that shows some statistical advantages specifically when looking at the low and high end of the methylation range^42^. CimpleG also receives a vector *y* ∈ {0, 1}^*p*^, which is 1 if observation *j* is of a cell type of interest and zero otherwise (others).

CimpleG characterises a DNAm site *i*, for the cell type of interest, by the difference in means (Eq. 1) and the sum of variances (Eq. 2).

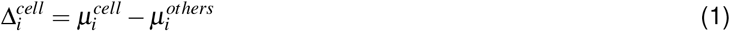

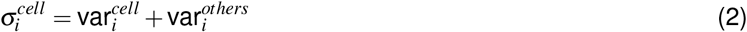

where 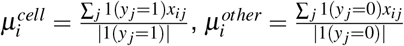 and 1 is an indicator function. Similarly, 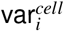 and 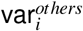 provides variance estimates for DNAm site *i*.

CimpleG first selects active features such that Δ_*i*_ is high and σ_*i*_ is low. For this, it ranks DNAm sites according to *a* with

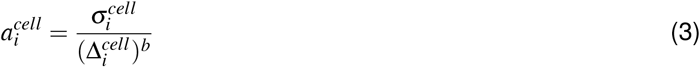

where *b* is a non-negative even constant integer that determines how much importance features with high difference in means should have. The larger *b* is, the larger the bias will be towards selecting DNAm sites with a higher difference in means Δ, regardless of the sum of variances σ. Of note, for *b* = 1, Eq. 3 is equivalent to a *t*-student statistic for the comparison of mean values of two groups. We set *b* to 2 as default. Finally, a quantile function *Q*(*p*) is used to generate a threshold below which sites with a value of *a* are selected. By default, we select 0.5% sites as active features.

CimpleG performs a balanced stratified K-fold cross-validation loop on the training data and uses the previous procedure to find the *f* active features for each fold. CimpleG sets *k* =10 as a default number of folds, unless the number of target class samples *n* < *k*, in which case *k* = *n*. To evaluate individual features as one-feature classifiers, it uses an area under the precision-recall curve (AUPR) procedure. This methodology is inspired by a feature selection method based on the area under the curve (AUC) proposed by Chen and Wasikowski^25^. In short, we evaluate the precision and recall for a linear classifier *x*_*i*_in regards to *y*. CimpleG adopts an AUPR curve instead of AUC as an accuracy metric, as it makes the procedure less affected by class imbalance^44^.

After evaluation of all folds, we compute a final metric by considering DNAm sites *i*, which minimises the score *a*_*i*_, while maximising the average AUPR values in training and validation (*AUPR*_*ti*_ and *AUPR*_*vi*_ respectively) and being active features (*f*_*i*_) in most folds, that is:

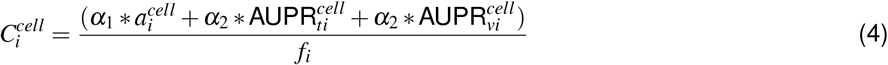

where *i* denotes a given feature. Finally, *α*_1_ and *α*_2_ are constants by which we control how strong the influence of *a* and the *AUPR* values will have on the final score. We set these as 0.8 and 0.2 respectively. We then use this score to rank all active features for a given cell type.

### Data

We have compiled and curated two human cell-type DNA methylation datasets that were measured with Illumina Infinium Human Methylation 450k and EPIC BeadChip arrays and deposited in the Gene Expression Omnibus (GEO)^45, 46^. Cell type information and other metadata were also obtained from GEO. Note that although some samples are shared between the somatic cells and leukocyte datasets, these were treated as independent datasets and therefore the samples were processed and treated as part of their respective dataset.

### Somatic Cells

We have searched for DNAm array data with an emphasis on purified, well-characterised, non-malignant cells in GEO. After manual curation, we had 576 samples, spanning over 14 different cell types, from 46 distinct studies (see Table 1 for overall statistics)^13^. We provide a detailed sample sheet in Supplementary Table 2, which includes GEO sample IDs (GSMs), GEO series/studies IDs (GSEs), cell type and assigned dataset (train/test).

### Leukocyte Cells

As with the previous dataset, we have searched for DNAm array data in GEO, however here we focused on purified leukocytes. After manual curation, we had 365 samples, for eight different cell types from 12 different studies (see Table 2 for overall statistics). We provide a detailed sample sheet in Supplementary Table 3, which includes GEO sample IDs (GSMs), GEO series/studies IDs (GSEs), cell-type and train/test dataset assignment.

### DNA methylation data and quality control

Both datasets were independently pre-processed. Raw data (.IDAT files) was downloaded from GEO. If .IDAT files were not available, the tabular data with probe intensities was used. For datasets where IDAT files were available, the SeSAMe pipeline^21^ was used. Briefly, this includes, probe filtering regarding low mapping quality, single sample normalisation (ssNoob)^18^, nonlinear dye-bias correction and p-value with out-of-band array hybridisation masking/filtering. For samples where only intensity matrices were available, beta mixture quantile normalisation (BMIQ, wateRmelon R package v1.34.0^28^) for type II bias correction was used. After merging the data obtained from the .IDAT files and the tabular data with probe intensities, we remove samples for which over half of the probes are missing, followed by eliminating all probes that still have missing data. Furthermore, we only considered CpGs that were provided by the 450K and EPIC BeadChip platforms. With this procedure, the final datasets have 143,291 and 284,706 features for the Somatic and Leukocyte cells, respectively. This process filters out a large number of probes, however ensures that the probes kept have a higher mapping quality and should generalise better in unseen datasets.

#### Additional DNAm methods for classification

CimpleG also allows the user to train classifiers using a number of different machine-learning algorithms to generate complex methylation signatures. For this, we explore methods exposed by the tidymodels framework^30^ and provide a wrapper around five popular models with different levels of complexity (Elastic Net, Decision Trees, Random Forests, Boosted Trees and Neural Networks). Each method has a set of hyper-parameters that need to be tuned in order to produce proper results. We chose a sensible set (in regards to computational resources) of hyper-parameters to be tuned via grid-search in a stratified cross-validation (CV) loop. After the CV loop, we fit the models on the whole training dataset with the previously tuned hyper-parameters. Due to the fact that some methods did not cope with the original feature size (Decision Trees, Random Forests and Neural Networks) of DNA methylation data (>100.000 features), we had to apply a filter as a pre-training step in order to remove features that are highly sparse, features that show linear combinations between them and features that have large absolute correlations with other variables. These filters are applied as a pre-training step. The implementation of these methods is out of the scope of this paper but we describe them briefly here.

#### Elastic Net

Elastic Net^15^ is a regularised linear regression method that is able to perform feature selection in arbitrarily large datasets. It can also be used for two-class classification problems within a logistic regression framework. Elastic Net has a regularisation parameter *γ* that controls penalisation, where higher values indicate higher penalisation (lower number of features). Additionally, the mixture parameter *α* controls the type of regularisation, where if *α* = 0 we have a pure ridge regression model, and if *α* = 1 we have a lasso regression model. Here, we optimise both *α* and *γ* during cross-validation. We used the glmnet package^16^ as the underlying engine for these models.

#### Decision trees

Decision trees are simple models that are defined by a set of if/else statements creating a tree-based structure. These models are very easy to understand, but they tend to be inaccurate and unstable and also have several hyper-parameters to tune. We used the rpart package^47^ as the underlying engine for these models. Here, we consider the maximum tree-depth as the single parameter to be tuned during the CV loop, while most other parameters are set as default values. The exceptions are the engine-specific parameters “maxsurrogate” and “maxcompete” which were set to 2 and 1 respectively, in order to reduce execution times.

#### Random forests

Random forests is an ensemble based method that builds predictions by combining decision trees in a bagging fashion. In essence, random forests create a large ensemble of independent decision trees, which are based on a random and small selection of features, and then builds its predictions using a combination of the predictions of all the individual decision trees. We used the package ranger^48^ as the underlying engine for these models.

Here, we set the number of independent decision trees as the parameters to be tuned during the CV loop, while other parameters, except for “oob.error”, “respect.unordered.factors” and “importance”, were set to default. The aforementioned engine-specific parameters were set to “FALSE”, “order” and “impurity_corrected” respectively. This was in order to save on computation time, to better fit our experimental design (2-class classification problem) and to be able to measure feature importance after model training.

#### Boosted trees

Boosted trees are also models that are built on ensembles of decision trees. While boosted trees also typically create a large number of decision trees, the selection of samples and features are based on the predictive performance of the previously trained decision trees. To perform predictions all the decision trees in the ensemble are combined to produce a result. Here, we used the package xgboost^49^ as the underlying engine for boosted trees. Similarly to the random forests models, we set the number of trees in the ensemble as the parameter to be tuned during the CV loop. To improve computation times and to better fit our experimental design, we set the parameter “mtry” to 100. Furthermore, we set the engine-specific parameters “objective” to “binary:logistic”, “eval_metric” to “aucpr” and finally, “maximize” to “TRUE”. Other parameters were kept as default as per the tidymodels interface.

#### Neural networks

Neural networks are models that work with the concept of combining layers of interconnected computational neurons (perceptrons) to produce a prediction. Here we focus on a simple, feed-forward neural network with three layers, the input layer, a single hidden layer and the output layer. We used the nnet package^50^ as the underlying engine for our networks. In our wrapper implementation, we set the number of hidden units (perceptrons in the hidden layer) and the complexity penalty parameter as the parameters to be tuned during the CV loop. To have a sensible computational footprint, we kept the epochs (number of training iterations for each network) to 100, as default. However due to the nature of our data (high-dimensionality, even after filtering), we had to set the engine-specific parameter “MaxNWts”, which controls the maximum number of weights in the model, to one million, otherwise, the models would not fit. This parameter is necessary but makes the fitting process more computationally expensive.

### Experimental design

The overall design is shown in Supplementary Fig. S2. First, we divided the data into a train and a test set. The train set was used to train and fit the models within a cross-validation setup while the test set was used as an independent assessment of the models after training concluded. The split in the data was manually curated. To avoid bias, we had to ensure that (1) samples from the same study were exclusively in the train set or the test set and (2) whenever possible given (1), the proportion of samples for a given cell type across the different cell types was kept stable between the train and test sets so that these were as balanced as possible. Normalisation was performed independently for the train and test data as previously described. Since some algorithms (neural networks, decision trees and random forests) cannot cope with the high dimensionality of the data, CimpleG enforces, during training but before the cross-validation step, a correlation, a co-linearity and a variance-based filter. Therefore, for these three algorithms, the universe of features is reduced to 360 and 402 CpGs for the somatic and leukocytes datasets respectively.

Next, we performed model training for every evaluated approach and for every relevant cell type within the somatic cells and leukocyte datasets (Table 1 and 2, respectively). Note that, each model is trained independently for each different target class. This means that, for example, we will have six different and trained random forest models for the leukocytes dataset (that has 6 different target classes). For model selection and parameter optimisation, we performed stratified cross-validation (10-fold) in the training dataset. The same folds were used for all evaluated models and cell types. See the text above for a description of the optimised parameters. Finally, we assess the performance of each method using the test dataset. The classification performance of each method is evaluated using the Area under the Precision-Recall curve. Other metrics considered for the evaluation and comparison of the models were the time of execution (which includes model optimization) and the number of selected features (if supported by the method). To quantitatively understand how each model ranks against the other, we computed the overall mean ranks (based on the AUPR) as well as the Friedman-Nemenyi post-hoc tests. With these, we can assess if different methods perform significantly better than others (Fig. 2).

### Deconvolution analysis

Deconvolution is often posed as a matrix factorization problem where

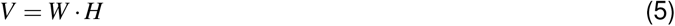

In this context, *V* would generally be the methylation level from different samples (the data we are trying to deconvolute), *W* would be the cell-type specific methylation values (our reference matrix) and finally, *H* would be the cell-type proportions. Therefore, we would like to find *H*, given that we have *V* and *W*. Given that all these matrices are positive, this problem can be solved with a non-negative least square algorithm (NNLS)^35^.

For our signatures, we generate *W* in two distinct ways depending on the type of model the signatures originated from. If the signatures were generated from one of the simple methods (CimpleG or brute-force), then we take their DNAm site selection and build a reference matrix using the average DNAm values of each CpG per class label. Otherwise, for each machine learning method, we build the reference matrix by using the average classification score per classifier, per cell type. To perform cellular deconvolution on the matrix *V*, we again use two different approaches. For the simple methods, we take the methylation level of the DNAm signature sites. However, for the complex methods, we use the classification score of each classifier for the different samples. Taken together, the *W* and *V* matrices are fed to the NNLS algorithm to reconstruct *H*.

To quantitatively evaluate the performance of different methods for deconvolution we used the root mean squared error (RMSE) and r-squared (*R*^2^)^51, 52^ as performance measures on 12 artificially mixed samples with known proportions from a recent study^36^. These artificial mixtures consist of DNAm profiles from purified leukocytes, that are combined with known proportions to simulate blood^36^. These samples have also been analyzed with flow-cytometry, making these an ideal dataset to quantitatively evaluate deconvolution. We adapted the cell-proportion labels to reflect our trained models and signatures. Namely, we condensed their multiple subgroupings of CD4+ T-cells, CD8+ T-cells and B cells into just these labels. Further, we merged their neutrophils, eosinophils and basophils labels into a general granulocytes group.

## Data and software availability

CimpleG’s source code can be found at github.com/CostaLab/CimpleG and its website with tutorials, examples, documentation and code used for this manuscript can be found at costalab.github.io/CimpleG.

## Acknowledgements

This work was funded by grants of the Interdisciplinary Center for Clinical Research (IZKF) Aachen, RWTH Aachen University Medical School, Aachen, Germany (to W.W. and I.C.); by the Deutsche Forschungsgemeinschaft (DFG-GE 2811/3 to I.C, WA 1706/12-1 to W.W. and WA1706/14-1 to W.W.) and by the Bundesministerium für Bildung und Forschung (BMBF; e:Med Consortia Fibromap to I.C. and VIP+ Epi-Blood-Count to W.W.), and the ForTra gGmbH für Forschungstransfer der Else Kröner-Fresenius-Stiftung (to W.W.).

## Author contributions

T.M. performed the computational analysis, wrote the source code for CimpleG and the initial draft of the manuscript.M.S compiled and curated the methylation data sets, provided suggestions and insight for the development of CimpleG and pre-processing of the datasets. M.S. and M.E. helped debugging and improving initial versions of the software. M.E. and I.C. annotated the CpG signatures. I.C. and W.W. designed and supervised the study. T.M., M.S., I.C. and W.W. conceptualised the project. All authors revised the manuscript and approved the final version.

## Competing interests

The authors declare no competing interests.

## Notes

https://costalab.github.io/CimpleG/

### Competing Interest Statement

The authors have declared no competing interest.

