## Supplementary Figures for "CimpleG: Finding simple CpG methylation signatures"

### Supplementary material

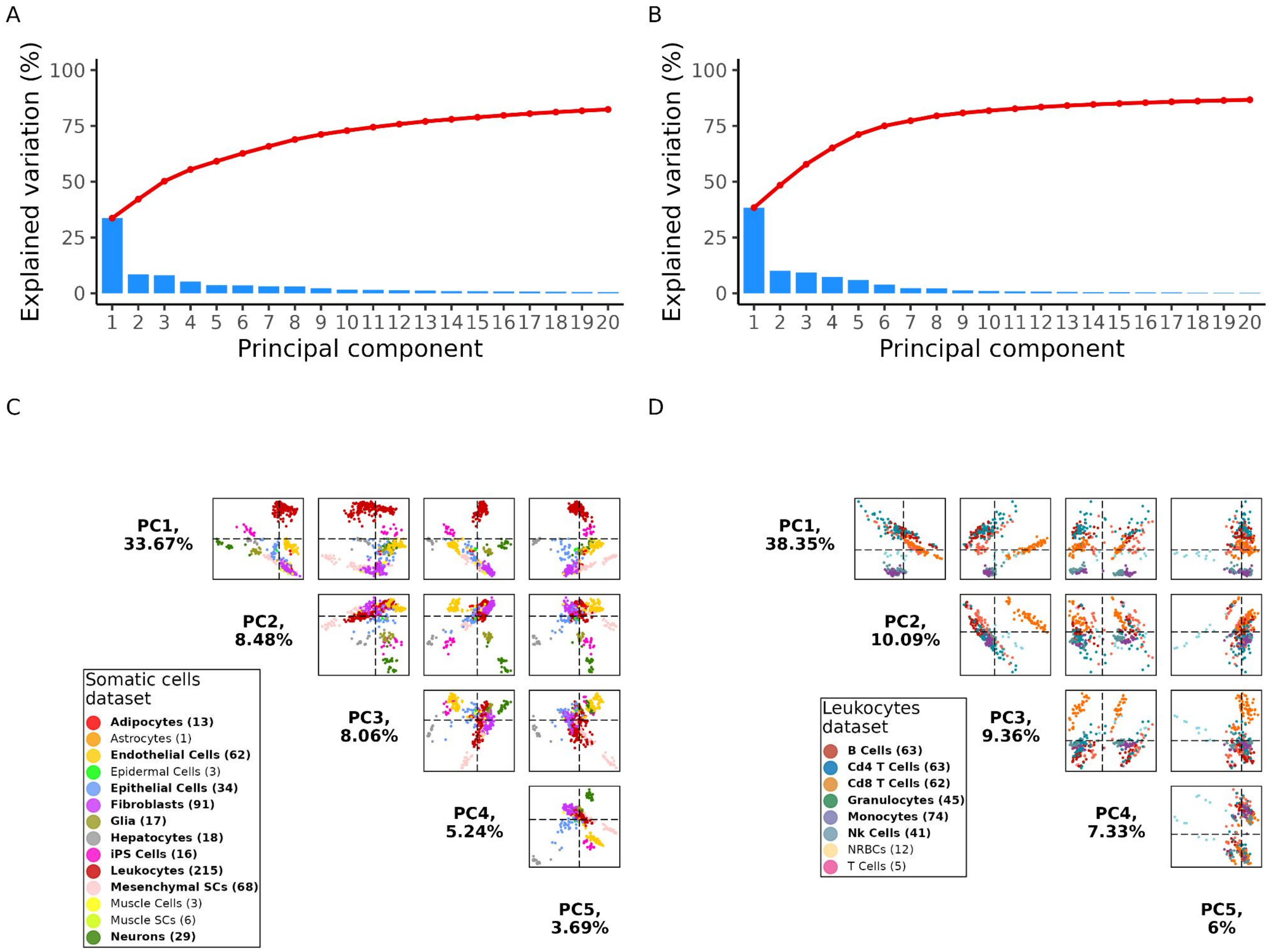


**Fig. S1.** Principal component analysis of the datasets used. (A-B) Scree plots from the PCA analysis for the somatic cells dataset and the leukocytes dataset respectively; (C-D) PCA pair-plots with the first 5 Principal Components for the somatic cells dataset and the leukocytes dataset respectively. The separation of individual cell types varies across the different principal components.

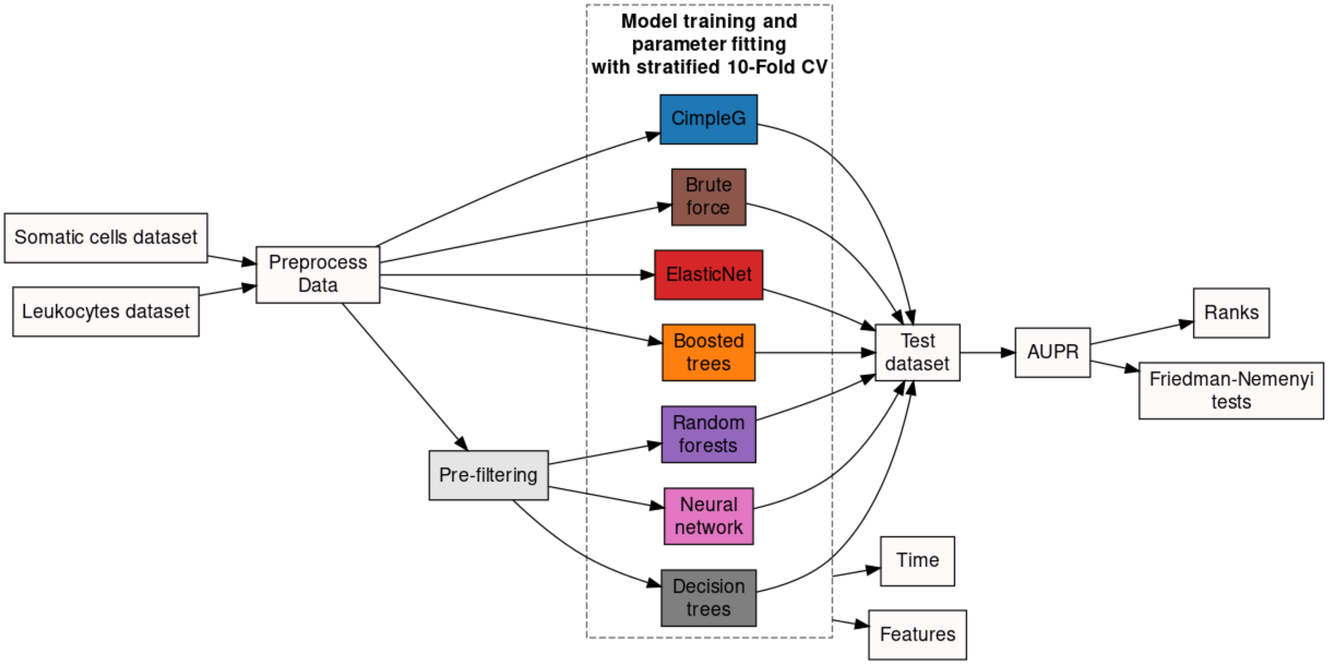


**Fig. S2.** Schematic representation of the experimental design used for benchmarking.

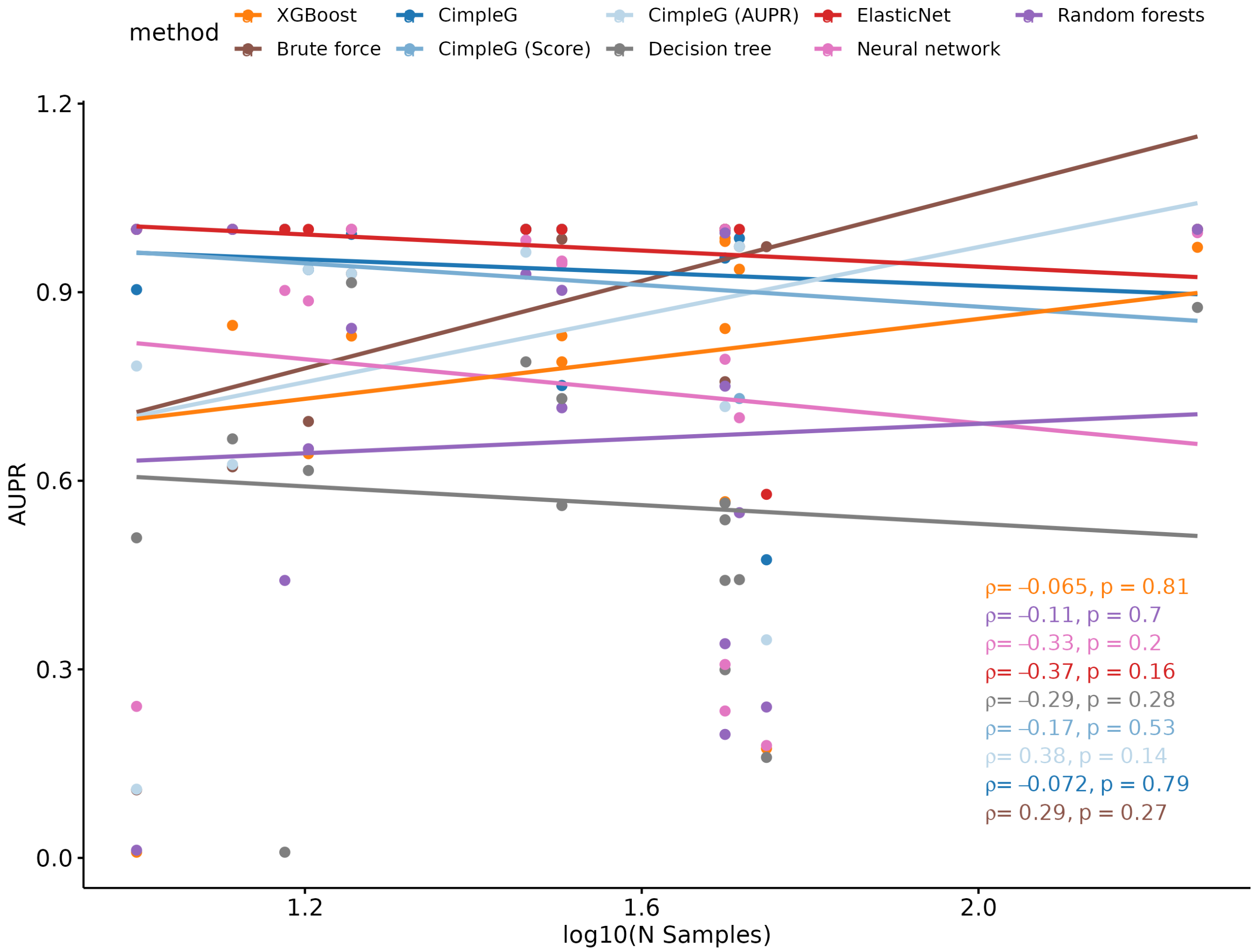


**Fig. S3.** Relationship between the Area Under the Precision–Recall curve (AUPR) and the number of samples for the target class (in the log10 scale) for all evaluated methods and datasets. Lines represent linear fits, which evaluate if the AUPR of distinct methods is related to the sample size of the target class. Statistics shown are Spearman's correlation coefficient rho and the associated p-value.

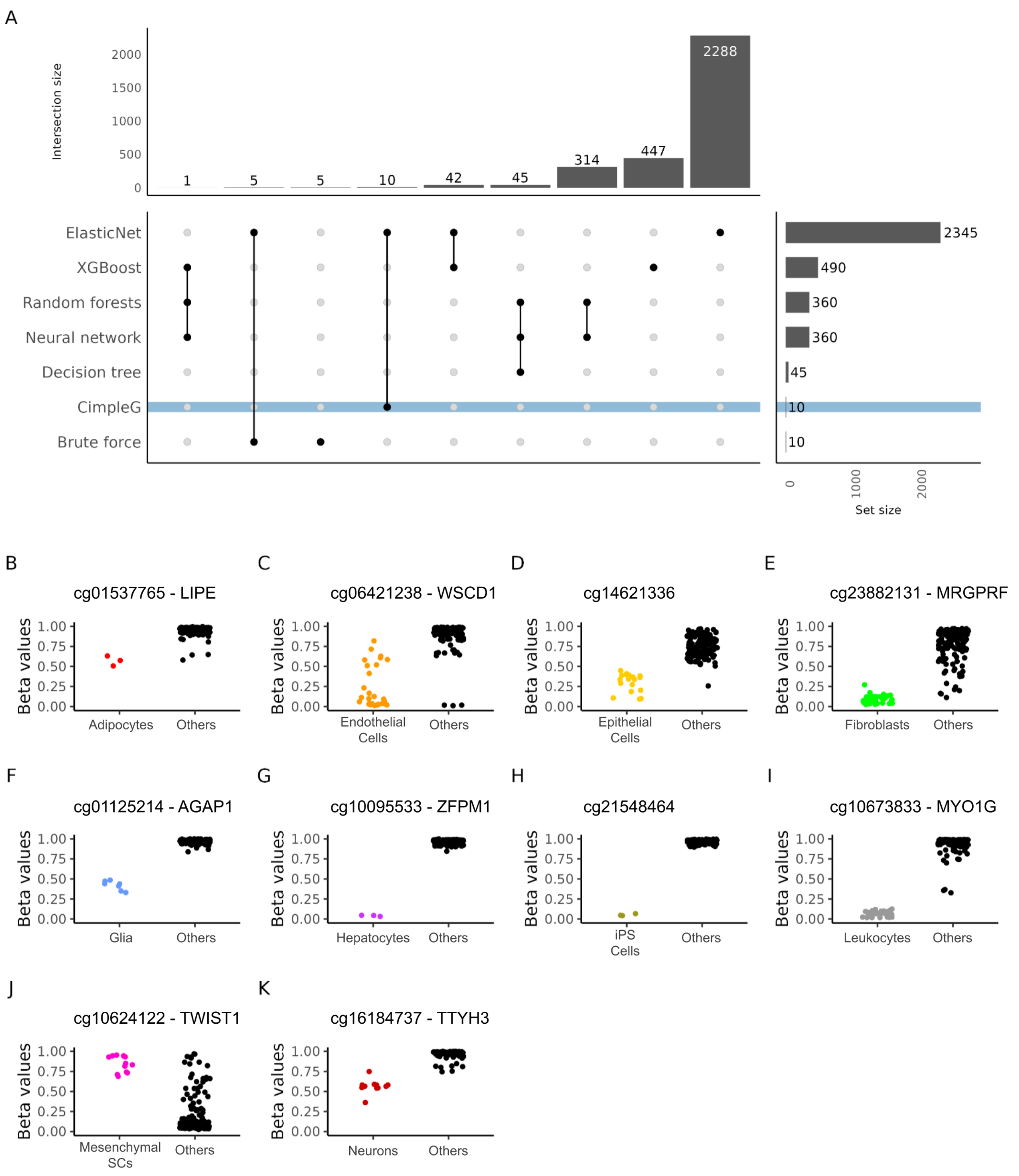


**Fig. S4.** Somatic cell signatures. (A) Upset plot showing the total number of selected DNAm sites per method (y-axis) and how these DNAm sites intersect for distinct combinations of method (x-axis) for the somatic cells dataset. Connected dots in a column indicate the combination of methods. (B-K) Beta values (y-axis) of CpG sites selected by CimpleG on the test data . The colour of the points corresponds to the target cell type, while points in black correspond to the cell types that are not the targets for that signature.

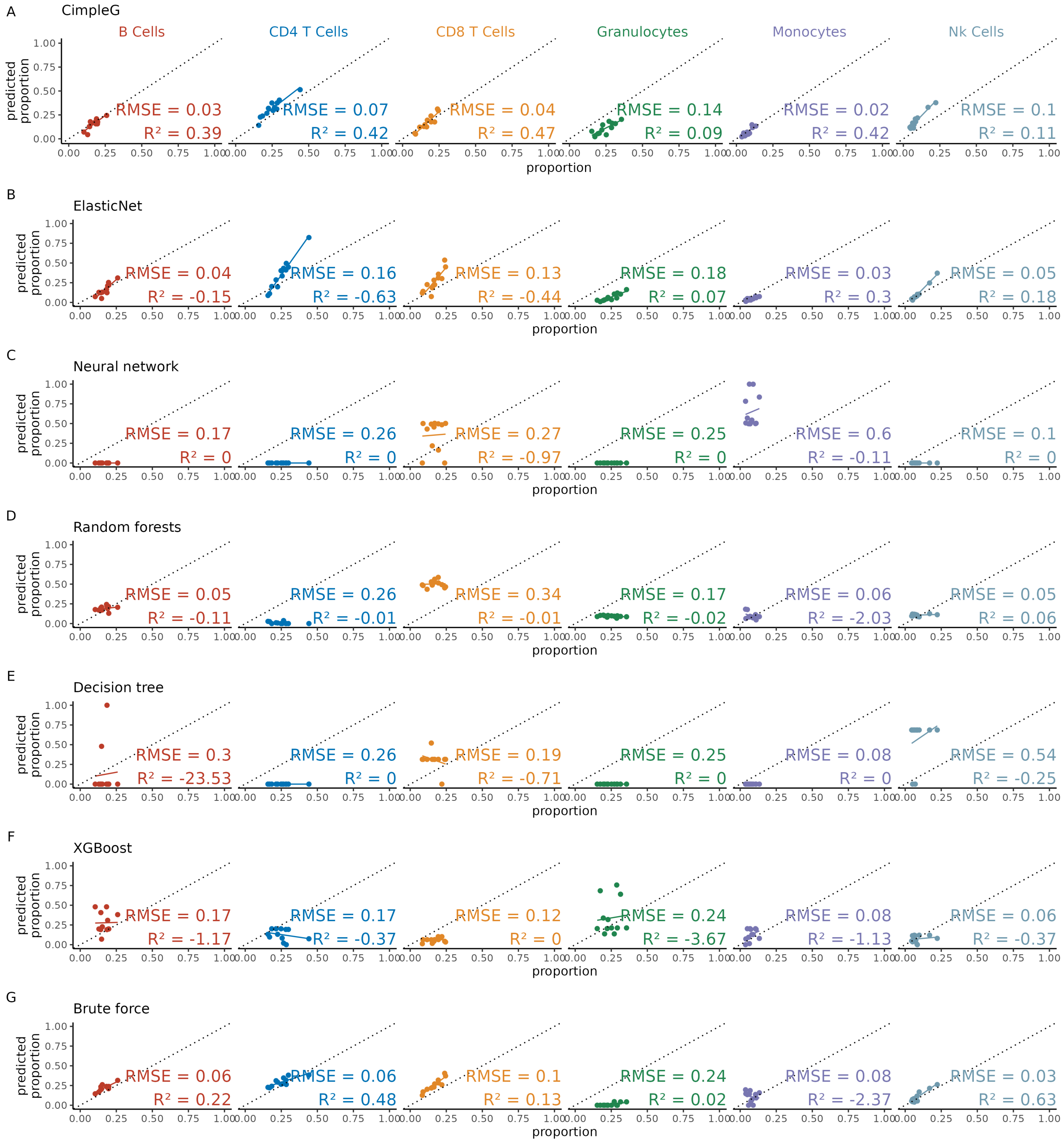


**Fig. S5.** The predicted deconvolution proportion (y-axis) vs real proportion (x-axis) for all leukocyte mixtures for all the trained methods: CimpleG (A); ElasticNet (B); Neural network (C); Random forests (D); Decision trees (E); Boosted trees (F) and Brute force (G). Predictive accuracy statistics, RMSE and $R^2$, are shown for each trained classifier.
